# Nanopore sequencing-based measurement of paramyxovirus RNA editing reveals virus-specific differences in editing efficiency of mRNA, antigenome and genome

**DOI:** 10.1101/2025.02.16.638481

**Authors:** Yusuke Masuda, Tofazzal Md. Rakib, Lipi Akter, Keisuke Nakagawa, Kiyotada Naitou, Akatsuki Saito, Ryoji Yamaguchi, Yusuke Matsumoto

## Abstract

Paramyxovirus polymerase recognizes an RNA editing signal on the viral genome and transcribes mRNA in which guanine nucleotides are inserted in a template-independent manner. This enables the synthesis of multiple proteins from a single gene, which is important for viral growth. We developed a method to quantify RNA editing efficiency using Oxford Nanopore Technologies’ MinION platform. We performed sequence analysis of reverse transcription (RT)-PCR amplicons with the RNA editing sites in cells infected with Sendai virus (SeV) and canine distemper virus (CDV). By modifying RT primers, we simultaneously assessed RNA editing efficiency in mRNA, antigenome and genome. We observed distinct differences in mRNA editing efficiency between SeV and CDV. Notably, while RNA editing in SeV is confined to mRNA, in CDV it is also observed in antigenome/genome. (Anti)genomes harboring extra nucleotides may deviate from a multiple-of-six sequence, suggesting that RNAs not following the “Rule of Six” are produced in CDV-infected cells.

**Highlights:**

- A method was established to quantify RNA editing efficiency in RNAs from paramyxovirus-infected cells using the MinION platform.
- RNA editing efficiency in mRNA differs between Sendai virus and canine distemper virus.
- Although RNA editing is confined to mRNA during Sendai virus infection, it is also observed in the antigenome and genome during canine distemper virus infection.

## Introduction

Paramyxoviruses are divided into three major subfamilies: *Orthoparamyxovirinae, Rubulavirinae* and *Avulavirinae* (Duprex and Dutch, 2023). The *Orthoparamyxovirinae* comprises the genera *Respirovirus, Morbillivirus* and *Henipavirus*, which include numerous viruses that cause infectious diseases in humans and various animals. Sendai virus (SeV), in the genus *Respirovirus*, is known to infect rodents and has significantly contributed to the understanding of growth mechanisms of paramyxoviruses. Canine distemper virus (CDV), in the genus *Morbillivirus*, infects canids and causes systemic diseases.

All paramyxoviruses share a common mechanism for genome replication. In SeV, the negative-sense single-stranded RNA genome encodes six genes arranged in the order 3′–N–P/V– M–F–HN–L–5′, with each gene separated by a gene junction. This negative-sense genome is completely encapsidated by the nucleoprotein (N) and is recognized as a template by the RNA-dependent RNA polymerase (RdRp) complex, which consists of the large protein (L) and an accessory factor, the phosphoprotein (P). The RdRp complex sequentially transcribes mRNA using the negative-sense genome as the template and also replicates a positive-sense antigenome that later serves as the template for progeny negative-sense genomes. Moreover, paramyxoviruses adhere to the so-called “Rule of Six,” meaning that the genome must consist of a number of nucleotides that is a multiple of six for replication to occur (Calain and Roux, 1993), and the N protein encapsidates the genome and antigenome in precise increments of six nucleotides (Alayyoubi et al., 2015; Gutsche et al., 2015; Luo et al., 2020).

Paramyxoviruses also exhibit characteristic RNA editing within the P/V gene region. RNA editing is a phenomenon in which the RdRp, upon recognizing a specific editing site on the genome, inserts guanine nucleotide(s) into the transcript in a template-independent manner with a certain probability (Cattaneo et al., 1989; Douglas et al., 2021; Vidal et al., 1990a). This insertion causes a shift in the mRNA open reading frame, thereby altering the protein that is subsequently translated. In *Orthoparamyxovirinae* subfamily, the P protein is translated when the RNA editing does not occur; however, a one-nucleotide shift results in the production of the V protein. The P protein is an essential polymerase cofactor for viral replication, while the V protein is an accessory protein that interacts with multiple host cellular proteins and acts as an interferon antagonist (Chambers and Takimoto, 2009; Wang et al., 2022). Thus, RNA editing determines the balance of P and V protein expression and has a significant impact on viral replication. Accurate measurement of RNA editing efficiency is important for elucidating the mechanisms of viral propagation. Although primer extension-based assay or cDNA sequencing for cloned mRNAs have been predominantly used to measure RNA editing frequency in viral mRNAs (Haas et al., 1995; Iseni et al., 2002; Vidal et al., 1990a), it has limitations in precisely quantifying the various RNA editing products.

In this study, we report a method for quantifying the RNA editing rate of mRNA produced in cells infected with paramyxoviruses, by combining reverse transcription-PCR with the MinION platform from Oxford Nanopore Technologies. By optimizing the primers used for reverse transcription, we determined the RNA editing efficiency in mRNA, antigenome and genome. This approach not only revealed differences in RNA editing efficiency in mRNA between SeV and CDV—viruses belonging to different paramyxovirus genera—but also demonstrated that, in CDV, RNA editing occurs in the antigenome, resulting in the emergence of edited genomes.

## Materials and Methods

### Cells and viruses

Vero cells (Japanese Collection of Research Bioresources Cell Bank [JCRB], Ibaraki, Japan, Cat# JCRB0111) and Vero cells stably expressing canine SLAM (Vero.DogSLAMtag; Vero-DST) (JCRB, Cat# JCRB1904) (Seki et al., 2003) were cultured in Dulbecco Modified Eagle’s Medium (DMEM) with 10% fetal calf serum (FCS) and penicillin/streptomycin. Cells were cultured at 37°C in 5% CO_2_. Sendai virus strain 52 was obtained through BEI Resources, NIAID, NIH: NR-3227. Canine distemper virus strain Vn99 is a field isolate as shown previously (Lan et al., 2009). Virus stocks for SeV and CDV were prepared using Vero and Vero-DST cells, respectively. The stock virus titers were calculated by the tissue culture infectious dose 50% (TCID_50_) determined using Vero and Vero-DST cells cultured in 96 well plates, respectively.

### Virus infection

Vero and Vero-DST cells (1 × 10^5^ cells/well) were cultured in 12 well plates, then infected with SeV and CDV at a multiplicity of infection (MOI) of 0.01 for 48 hours, respectively. For CDV, the infected cells were cultured in DMEM; for SeV, the infected cells were cultured in DMEM with 1.2% TrypLE select reagent (Thermo Fisher Scientific, Waltham, MA, USA).

### Reverse transcription-PCR

cDNA synthesis was carried out using PrimeScript II (Thermo Fisher Scientific) according to the manufacturer’s instructions. The primers used for reverse transcription and PCR amplification are listed in Table 1. All cDNA samples were subjected to PCR in 50 µL reaction mixture containing PrimeSTAR MAX (Takara Bio, Shiga, Japan) according to manufacturer’s instructions. Thermal cycling conditions were as follows: an initial denaturation at 98 °C for 1 min, 30 cycles of 98 °C for 10 s, 56 °C for 5 s and 72°C for 10 s; followed by a final extension at 72 °C for 1 min. The PCR products were subjected to agarose gel electrophoresis and purified by using a FavorPrep GEL/PCR Purification Mini Kit (Favorgen Biotech Corporation, Ping Tung, Taiwan) according to manufacturer’s instructions.

**Table 1.**
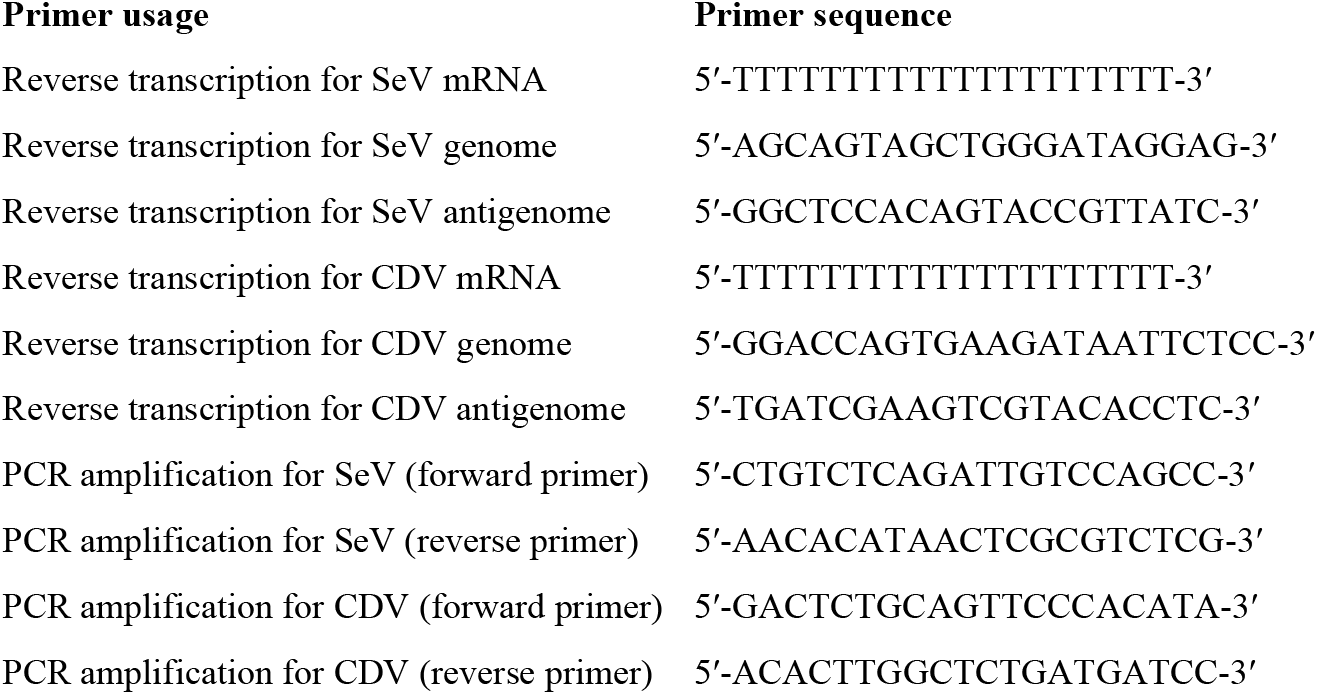
Primer sequences used for reverse transcription and PCR amplification.

### Nanopore sequencing

Oxford Nanopore sequencing was performed using the MinION portable nanopore sequencer with a single Flow Cell (R10.4.1) Nanopore FLO-MIN114. Library preparation was performed according to the protocol for the Native Barcoding Kit 24 v14 (SQK-NBD114.24, Oxford Nanopore Technologies, Oxford, UK). Library preparation involved end repair, adapter ligation, and purification steps to ensure high-quality sequencing-ready DNA. The prepared libraries were quantified using a Qubit fluorometer and loaded onto the flow cell. Sequencing was conducted using MinKNOW software to control the run parameters, monitor real-time data acquisition, and ensure optimal pore occupancy. The quality-check (QC) passed sequences were retrieved as FASTQ format and used to analyze the RNA editing quantification. The R script for RNA editing counting is available at https://raw.githubusercontent.com/ymatsu51/RNAediting/main/RNAediting.

### Data availability

The dataset of amplicon sequencing is available in the DNA Data Bank of Japan (DDBJ) under accession code (*to be assigned*).

## Results

To evaluate the RNA editing efficiency in SeV-infected cells, the virus was inoculated into Vero cells, and total RNA was extracted 48 hours post-infection. SeV produces three types of RNA in the infected cells: positive-sense mRNA, positive-sense antigenome, and negative-sense genome (Fig. 1). To analyze the sequence of the RNA editing region present in each of these RNAs, reverse transcription was performed on the segments containing the region. Since paramyxoviral mRNA is polyadenylated, an oligo-dT20 primer was used to specifically reverse transcribe mRNA (Fig. 1A). In contrast, although the genome and its complementary antigenome contain gene junction regions between each gene, mRNA lacks these junctions. Therefore, a reverse transcription primer was designed and positioned within the N gene in the appropriate orientation so as to reverse transcribe only the negative-sense genome (Fig. 1A). Similarly, for the antigenome, a primer was designed within the M gene in the correct orientation to specifically reverse transcribe the positive-sense antigenome (Fig. 1A). The reverse transcription products using these primers were used as templates to PCR-amplify, resulting in a 181-base-pair fragment within the P/V gene that contains the RNA editing region (Fig. 1B). An analogous procedure was applied to CDV-infected Vero-DST cells; an oligo-dT20 primer was used for mRNA, and CDV genome- and antigenome-specific reverse transcription primers were employed (Fig. 1A). The resulting reverse transcription products were then used as templates for PCR amplification of a 177-base-pair fragment (Fig. 1B).

**Figure 1.**
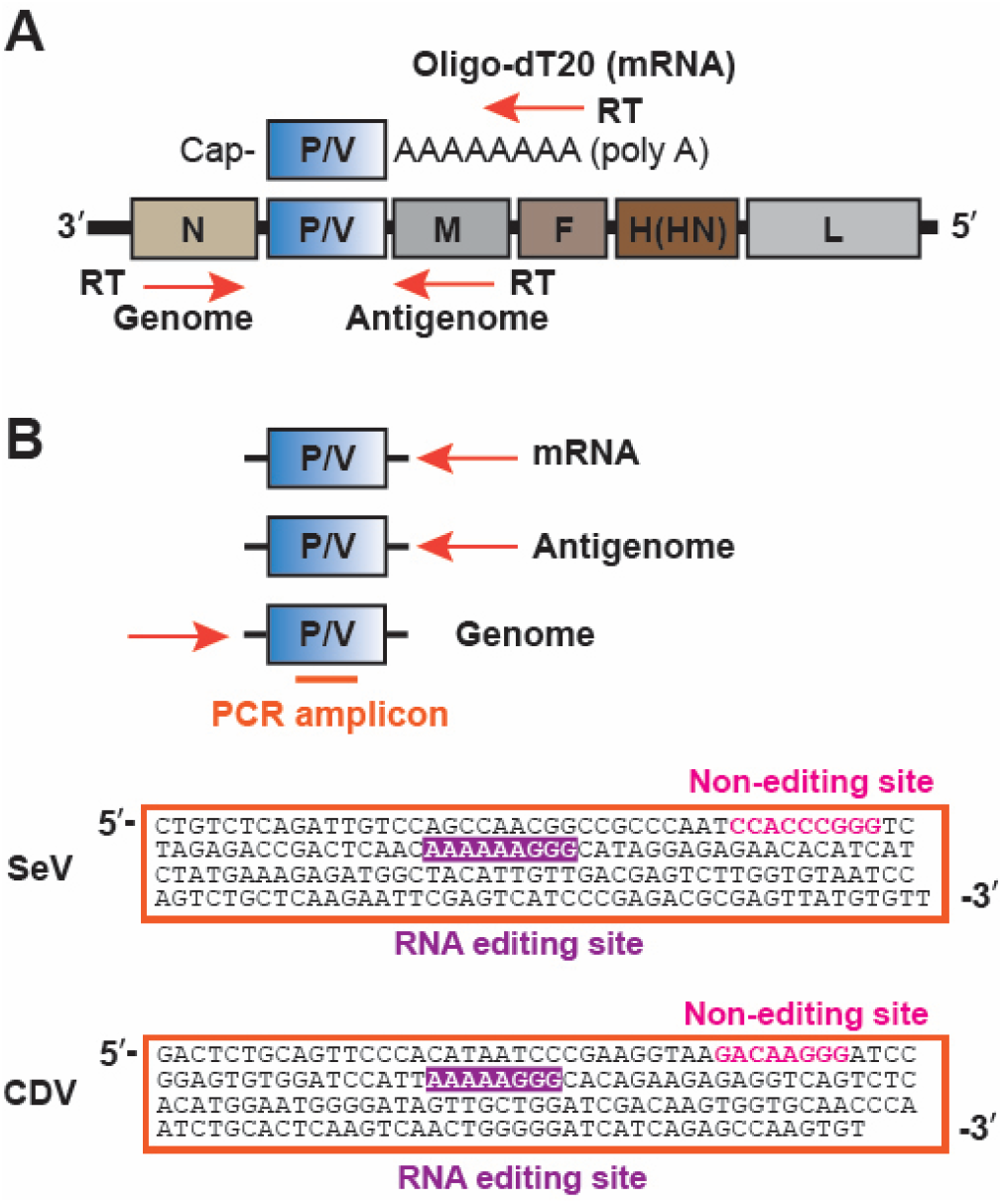
Amplicon sequencing of the editing region of SeV and CDV mRNA, antigenome and genome by the MinION platform. **(A)** A schematic diagram of the viral genome and the primers used for reverse transcription of each viral RNA in SeV- and CDV-infected cells. Three distinct primers were used to discriminate among mRNA, antigenome and genome. **(B)** The resulting cDNAs served as templates for PCR amplification of the P/V region containing the RNA editing site. The RNA editing sites, as well as the non-edited sites characterized by three G nucleotides for both SeV and CDV, are indicated. The PCR amplicons were subsequently sequenced using the MinION platform.

The resulting PCR products were subjected to sequence analysis using the Oxford Nanopore Technologies MinION platform. In the case of SeV, RNA editing occurs when the viral polymerase recognizes the RNA editing signal 3′-GUUGUUUUUUCCCGUA-5′ on the genome, leading to G nucleotide(s) being inserted in the transcript (e.g., resulting in sequences such as 5′-CAACAAAAAAGG**<u>G</u>**GCAU-3′) (Vidal et al., 1990b). Therefore, we counted the number of additional Gs in the corresponding region (5′-CAACAAAAAAGGGCAT-3′) of the obtained PCR product. The DNA sequence data in FASTQ format obtained from MinION were analyzed using an R script to count the number of sequences corresponding to 5′-CAACAAAAAAGGGCAT-3′ (G+0), 5′-CAACAAAAAAGG**<u>G</u>**GCAT-3′ (G+1), 5′-CAACAAAAAAGG**<u>GG</u>**GCAT-3′ (G+2), and 5′-CAACAAAAAAGG**<u>GGG</u>**GCAT-3′ (G+3) (Fig. 2A). The proportion of G+1 to G+3 was then calculated relative to the G+0 sequence count (set as 100%). The analysis revealed that in SeV mRNA the G+1 sequence accounted for approximately 30%, while G+2 and G+3 were present at less than 1% (Fig.2A). When the same analysis was performed on the antigenome and the genome, the proportions of G+1 to G+3 were found to be less than 1% in each case, suggesting that RNA editing in SeV occurs specifically in mRNA (Fig. 2A). To confirm that the addition of G nucleotides was confined to the RNA editing region, a separate region containing three consecutive G nucleotides—5′-AATCCACCCGGGTCTA-3′—present in the PCR products (Fig. 1B; Non-editing site) was similarly analyzed (Fig. 2B). In this case, the proportions of G+1 to G+3 were below 1% for mRNA, antigenome and genome (Fig. 2B). These results indicate that under the infection conditions of this study, RNA editing, manifesting as an approximately 30% occurrence of G+1 relative to G+0, occurs exclusively in SeV mRNA.

**Figure 2.**
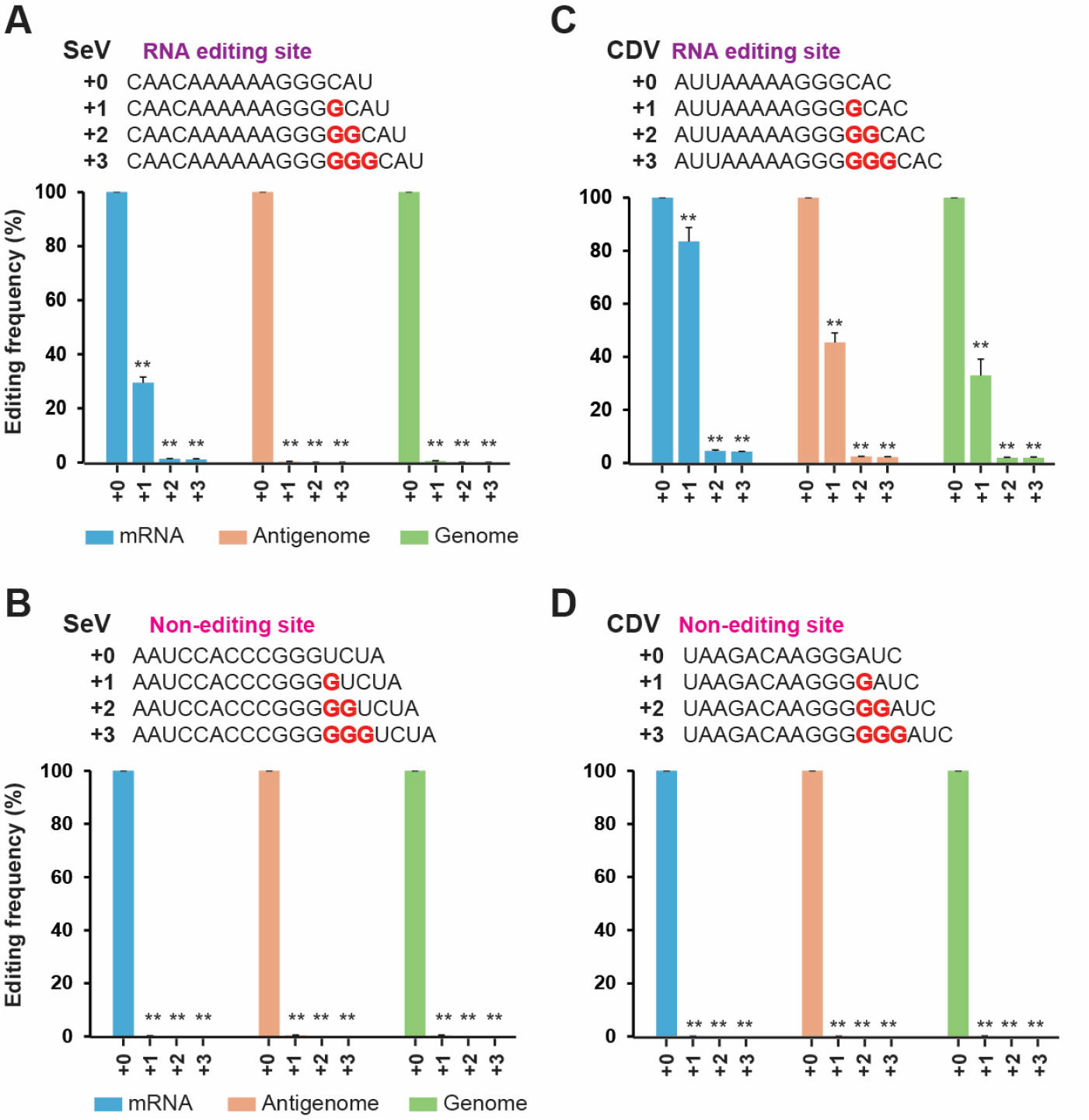
RNA editing efficiency for mRNA, antigenome and genome of CDV and SeV. Editing frequencies were calculated using an R script to analyze sequences obtained from the MinION platform for the RNA editing site of SeV **(A)**, the non-editing site of SeV **(B)**, the RNA editing site of CDV **(C)**, and the non-editing site of CDV **(D)**. The labels “+0” to “+3” indicate the number of additional G nucleotides present at the specified positions in the sequences. Editing frequencies are expressed as percentages, with the G+0 condition set at 100%. Data are presented as the mean ± standard deviation from three independent experiments. ^**^*p* < 0.01 (vs “+0”, a two-tailed unpaired Student’s *t*-test).

A similar sequencing analysis using the MinION platform was performed on CDV PCR products. For CDV, the polymerase recognizes the RNA editing signal 3′-UAAUUUUUCCCGUG-5′ on the genome, which results in G insertion(s) in the transcript (e.g., resulting in sequences such as 5′-AUUAAAAAGG**<u>G</u>**GCAC-3′) (Haas et al., 1995). We counted the number of additional Gs in the corresponding region (5′-ATTAAAAAGGGCAC-3′) of the obtained PCR product. An R script was used to count the numbers of sequences corresponding to 5′-ATTAAAAAGGGCAC-3′ (G+0), 5′-ATTAAAAAGG**<u>G</u>**GCAC-3′ (G+1), 5′-ATTAAAAAGG**<u>GG</u>**GCAC-3′ (G+2), and 5′-ATTAAAAAGG**<u>GGG</u>**GCAC-3′ (G+3), and the proportions of G+1 to G+3 were calculated relative to G+0 set at 100% (Fig. 2C). In CDV mRNA, G+1 was found to account for 84%, with G+2 and G+3 each constituting 4%. Furthermore, in the positive-sense antigenome, which is replicated using the negative-sense genome as the template, G+1 was observed at a high proportion of 45%, and in the negative-sense genome—which is replicated using the antigenome as a template—G+1 was present at 33% (Fig. 2C). In contrast, when a separate region with three consecutive G nucleotides (5′-TAAGACAAGGGATC-3′) that is not part of the RNA editing region (Fig. 1B; CDV) was analyzed, the proportions of G+1 to G+3 were less than 1% (Fig. 2D).

## Discussion

In this study, we established an experimental system that enables the facile quantification of RNA editing efficiency in the three types of RNA (mRNA, antigenome and genome) present in paramyxovirus-infected cells by employing RT-PCR products for sequencing using the Oxford Nanopore Technologies’ MinION platform. In SeV, RNA editing was observed exclusively in mRNA: approximately 30% of molecules exhibited the G+1 pattern when G+0 is set as 100% (Fig. 2A). Previously, a sequencing analysis of cloned cDNA containing the RNA editing region of SeV Z strain mRNA in infected BHK cells revealed that 99 clones exhibited the G+0 pattern, whereas 50 clones exhibited the G+1 pattern (Vidal et al., 1990a), corresponding to roughly 50% G+1 relative to G+0. In CDV, 84% of mRNA exhibited the G+1 pattern when G+0 equals 100% (Fig. 2C). A previous study employing two different CDV strains—the Onderstepoort and Yanaka strains—performed sequencing analysis of cloned cDNA containing the RNA editing region of mRNA in infected B95a cells. In both cases, when 20 clones exhibited the G+0 pattern, 8 clones exhibited the G+1 pattern (corresponding to 100% versus 40%) (Wakasa et al., 2000). It is suggested that RNA editing efficiency may vary depending on the strain of paramyxoviruses and the type of infected cells.

In SeV, RNA editing was not detected in either the antigenome or genome, indicating that RNA editing occurs exclusively in mRNA (Fig. 3A). In contrast, in CDV-infected cells, high-frequency RNA editing (G+1) was observed in both the antigenome and genome (Fig. 3B), suggesting that genomes and antigenomes containing one extra nucleotide are produced rather than those conforming to an exact multiple of six. The RNA editing signal in the CDV genome is thought to function as a cis-acting element not only during mRNA transcription but also during antigenome replication (albeit at lower frequencies—84% in mRNA and 45% in the antigenome). CDV antigenomes that retain the G+1 modification serve as templates for genome replication, and the G+1 detected in genome PCR products corresponds to a C+1 on the negative-strand genome (Fig. 3B; Genome). When C+0 is set as 100%, the occurrence of this modification was approximately 30% (Fig. 2C and 3B). This decrease from 45% in the antigenome to 30% in the genome is interpreted as resulting from the selective exclusion of 6n+1 antigenomes—which do not conform to the “Rule of Six”—during template recognition.

**Figure 3.**
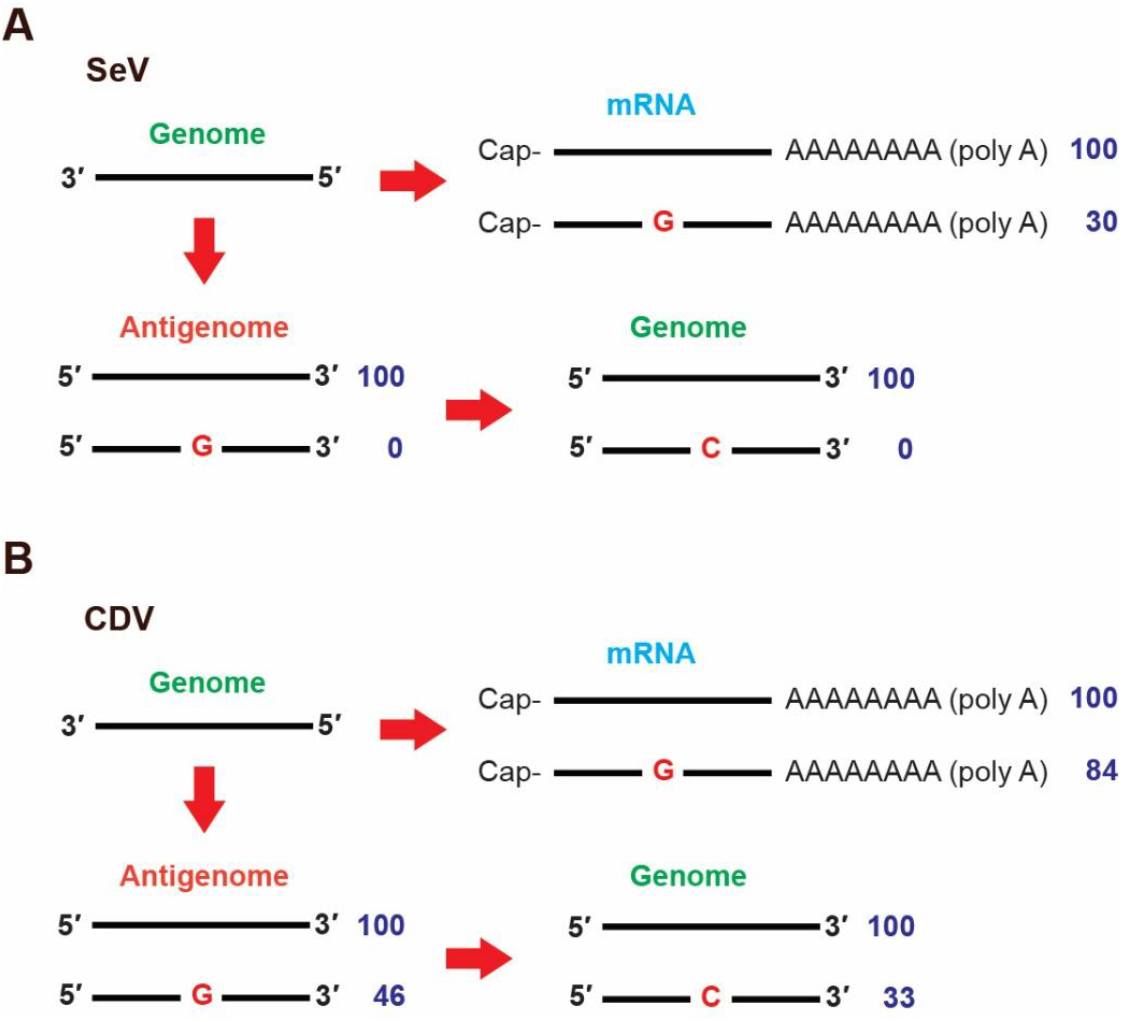
Schematic diagram of RNA editing in the mRNA, antigenome and genome. **(A)** Schematic diagram showing the viral RNA species in SeV-infected cells. Authentic mRNA editing results in 30% G+1 mRNA when G+0 is set to 100%. No RNA editing occurs during antigenome replication, and no edited genomes are detected. **(B)** Schematic diagram showing the viral RNA species in CDV-infected cells. Here, authentic mRNA editing results in 84% G+1 mRNA when G+0 is set to 100%. In contrast to SeV, the G+1 antigenome is detected at 46%, and the C+1 genome is detected at 33%.

The bipartite promoters play a critical role in enabling the paramyxovirus polymerase to discern whether a template is a multiple of six. Promoter Element 1 (PE1) is located at the 3′ end of the template, whereas PE2 is an internal promoter situated between the 73rd and 98th nucleotides from the 3′ end (Kolakofsky et al., 2021). The helical nucleocapsid of paramyxoviruses is formed by the precise binding of the N protein to RNA in each six nucleotides, and the functionality of PE2 depends on the proper positioning of the RNA sequence at the designated N-binding position. In SeV, the genomic PE2 sequence is “5′-AGGGU**<u>C</u>** UGGAA**<u>C</u>** CUGCU**<u>C</u>** UUCAGG-3′,” whereas in CDV it is “3′-UGAAC**<u>C</u>** CUAGC**<u>C</u>** UUGUU**<u>C</u>** UUUAAG-5′” (Ashida et al., 2024; Tapparel et al., 1998). In other words, within viruses of the *Orthoparamyxovirinae* subfamily, it is essential that the PE2 region consists of three consecutive repeats of “NNNNN**<u>C</u>**”— that is, a **<u>C</u>** residue must appear at three predetermined positions at every six-nucleotide interval in the N protein. If, during antigenome replication, an extra G is added in the RNA editing region, then the N proteins binding to the subsequently synthesized RNA will attach to sequences that are shifted by one nucleotide, causing the three consecutive C positions within PE2 to be misaligned. In such cases, the polymerase’s ability to recognize the template accurately is compromised, which is thought to be the factor preventing the viral polymerase from replicating templates that are not multiples of six. The observation that 45% of antigenomes in CDV exhibit the G+1 modification while only 30% of genomes display the corresponding C+1 is attributed to this selective mechanism. Nevertheless, the fact that as many as 30% of C+1 genomes remain suggests that the mechanism for selectively excluding templates that are not multiples of six may not operate as rigorously in CDV.

Although genomes that are not exact multiples of six may be produced in CDV-infected cells, CDV genomes isolated from the field are exact multiples of six (Ashida et al., 2024). This suggests that a selection mechanism may operate during the virion assembly process, ensuring that only genomes conforming to the Rule of Six are incorporated into viral particles.

Under the infection conditions used in this study, a higher mRNA editing efficiency was observed in CDV compared to SeV (SeV 30% vs. CDV 84%). This significant difference in mRNA editing efficiency may be directly reflected in the antigenome editing efficiency. The high mRNA editing efficiency of CDV might have led to editing of its antigenome as well. Thus, it is conceivable that, under conditions with inherently high mRNA editing efficiency, the antigenome in SeV might also undergo editing. Further detailed studies are needed to clarify this point.

## Conclusion

Our results showed that in SeV the RNA editing signal functions strictly during mRNA transcription, with no increase in nucleotide number observed in the RNA editing region of the antigenome and genome. In this case, only genomes that are exact multiples of six are present in SeV-infected cells, indicating that the Rule of Six is stringently maintained. In contrast, in CDV, the insertion of one nucleotide in the RNA editing region occurs frequently not only in mRNA but also in the antigenome and genome, suggesting that 6n+1-type genomes may appear in CDV-infected cells. These findings indicate that the strictness of the Rule of Six may vary among paramyxoviruses.

## Acknowledgements

The following reagent was obtained through BEI Resources, NIAID, NIH: Sendai Virus (formerly Parainfluenza Virus 1, Sendai), NR-3227. We thank Dr. Yusuke Yanagi (Kyushu University) for providing the Vero-DST cells.

